# Coral life history differences determine the refugium potential of a remote Caribbean reef

**DOI:** 10.1101/062869

**Authors:** Sarah W. Davies, Marie E. Strader, Johnathan T. Kool, Carly D. Kenkel, Mikhail V. Matz

**Affiliations:** Department of Integrative Biology, The University of Texas at Austin, 1 University Station C0990, Austin, TX 78712, USA; Department of Marine Sciences, The University of North Carolina at Chapel Hill, 123 South Road, Chapel Hill, NC 27599, USA; Geoscience Australia, Cnr Jerrabomberra Ave and Hindmarsh Drive, Symonston, ACT, 2609, Australia; Australian Institute of Marine Science, PMB 3, Townsville MC, Townsville, QLD, 4810, Australia

**Author notes:** These authors contributed equally to this work (Sarah W. Davies, Marie E. Strader).

## Abstract

Remote populations can influence connectivity and may serve as refugia from climate change. Here, we investigated two reef-building corals (*Pseudodiploria strigosa* and *Orbicella franksi*) from the Flower Garden Banks (FGB) – the most isolated, high-latitude Caribbean reef system that retains high coral cover. We characterized coral size frequency distributions, quantified larval mortality rates and onset of competence, estimated larval production, and created detailed biophysical models incorporating these parameters to evaluate source-sink dynamics from 2009 to 2012. Mortality rates were similar across species but competency differed dramatically: *P. strigosa* was capable of metamorphosis within 2.5 days post fertilization (dpf), while *O. franksi* were not competent until >20dpf and remained competent up to 120dpf Despite these differences, models demonstrated that larvae of both types were similarly successful in reseeding the FGB. Nevertheless, corals with shorter pelagic larval durations (PLD), such as *P. strigosa*, were highly isolated from the rest of the Caribbean, while long PLD corals, such as *O. franksi*, could export larvae to distant northern Caribbean reefs. These results suggest that FGB coral populations are self-sustaining and highlight the potential of long PLD corals, such as endangered *Orbicella*, to act as larval sources for other degraded Caribbean reefs.

## Introduction

Caribbean reefs have experienced some of the most dramatic coral declines over the last few decades^1^, however the Flower Garden Banks (FGB) – a system of two very unusual reefs located 185 km south of the Texas-Louisiana border in the Gulf of Mexico-appear to be the exception. The FGB is populated by only 24 species of reef-building corals^2^ but average coral cover exceeds 56%^3^, five times that of the general Caribbean average^1^. The FGB is one of the northern-most coral reefs in the Caribbean and is highly isolated from other reefs: the nearest neighboring reefs are hundreds of kilometers away along the coast of Tampico, Mexico (645 km) and the Yucatan peninsula (600 km)^4^ (Fig 1A). The FGB’s isolation, buffering from increased sea surface temperatures due to its high-latitude location, low degradation and high coral cover make it the ideal potential refugium from climate change for Caribbean corals. However, in order to be a good refugium the FGB must meet three requirements: first, FGB coral populations must be self-seeding (i.e., not requiring larval input from elsewhere to sustain populations), and second, coral larvae originating from the FGB must be capable of emigrating and surviving at other Caribbean reefs. Lastly, these reefs must be relatively resilient to recurrent disturbances such as bleaching.

**Figure 1.**
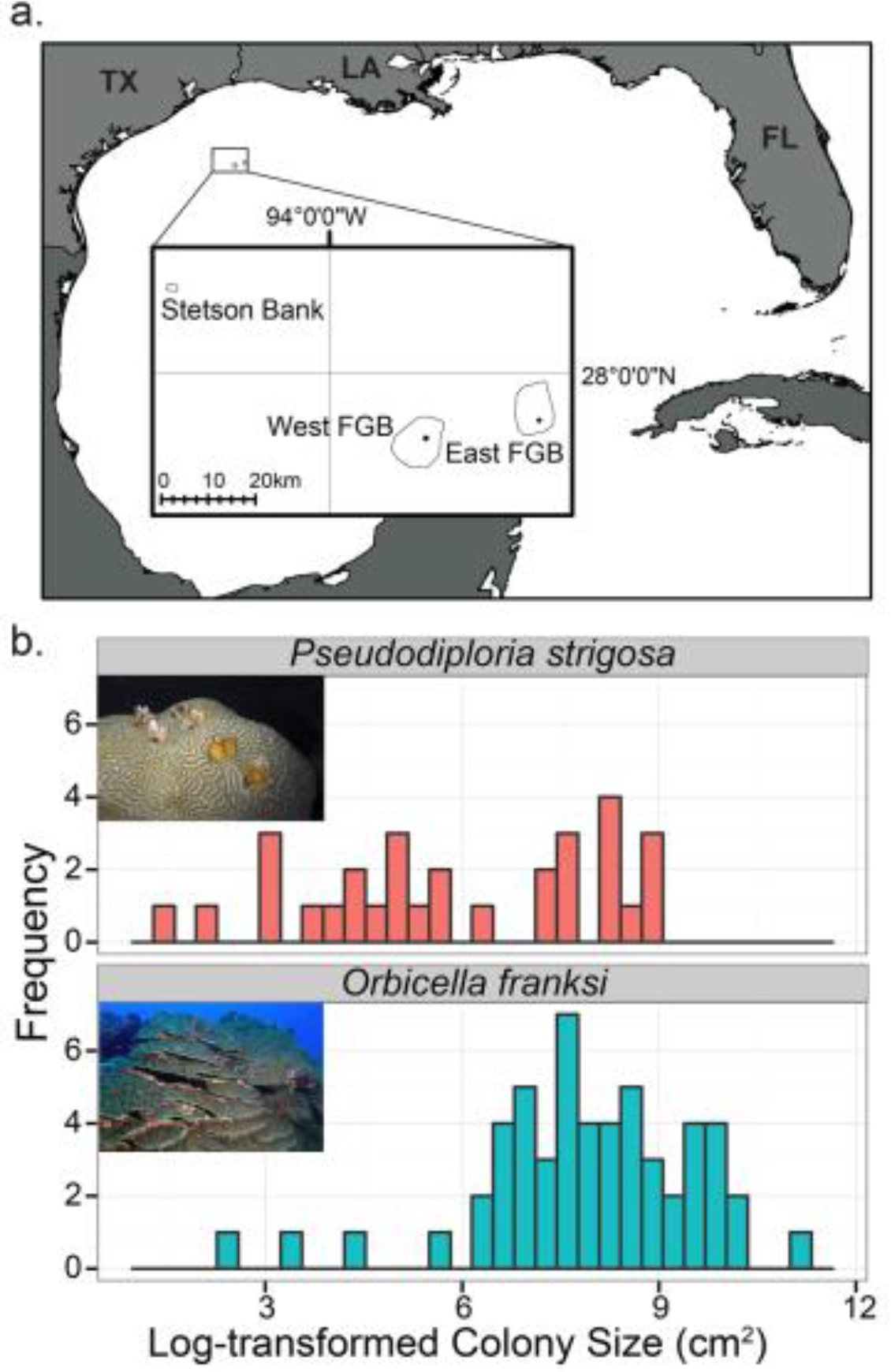
Size frequency data of *Orbicella franksi* and *Pseudodiploria strigosa* from the Flower Garden Banks (FGB) in 2012. A. Locations of surveys conducted at East FGB (mooring buoy 2) and West FGB (mooring buoy 2) (black dots). B. Size-frequency distributions for *P. strigosa* and *O. franksi* for both the East and West FGB resulting from transects performed in August 2012 with photo insets of each species.

Corals have bipartite life cycles with dispersive pelagic larvae, and a sedentary adult stage. Connectivity across distant populations is therefore dependent on the successful exchange of these pelagic larvae along ocean currents^5^. Intra and interspecies differences in biological traits can influence the scale of dispersal^6–8^. Larvae of some species have limited connectivity and only disperse meters from their parents^9^, while other species are highly genetically connected across distant reefs separated by thousands of kilometers^8,10,11^. The two key life history traits expected to influence connectivity are developmental time required before a larva becomes competent to settle, referred to as pelagic larval duration (PLD), and the larval mortality rate ^12,13^. In addition to life history traits, oceanic currents are strong drivers of larval connectivity and as global climates continue to warm, currents are predicted to dramatically shift, potentially altering the connectivity patterns of many marine species^14–16^. In order to fully understand population connectivity of reef-building corals it is essential to understand how ocean currents and life history traits interact to enhance or limit larval dispersal.

In this study, we aimed to estimate larval retention and export for the FGB. We used a biophysical model based on regional currents during the weeks following coral mass-spawning events across specific years (2009-2012) that also incorporated experimentally measured larval life history traits for two of the most dominant reef-building corals on the FGB: *Pseudodiploria strigosa* and *Orbicella franksi*. We find that our focal species represent opposite extremes in the timing of competency onset, which, unexpectedly, had little effect on their potential to reseed the FGB. However, only the species with delayed competency, *O. franksi*, had the potential for larval export to other Caribbean reefs.

## Results

### Size-frequencies and larval traits of <u>Psuedodiploria strigosa</u> and <u>Orbicella franksi</u> at FGB

Surveys of *O. franksi* and *P. strigosa* revealed similar size frequency ranges across the sampled reefs, however *P. strigosa* colonies were consistently in smaller size classes than *O. franksi* (Fig 1B; Wilcoxon sum rank test, P=2E-04). There was very little difference in larval mortality rates between *P. strigosa* and *Orbicella faveolata* (exponent powers of −0.022 and −0.019, respectively, Fig 2A), so mortality rates of both corals were modeled with the exponent power −0.02. The timing of competency onset varied dramatically between *P. strigosa* and *O. franksi. P. strigosa* exhibited competence on the very first trial, 2.5 dpf, and maintained competence at least until 8 dpf, at which point no swimming larvae remained in the cultures due to spontaneous metamorphosis (Fig 2B). In contrast, *O. franksi* became competent as late as 21 dpf and remained fully capable of metamorphosis at least until 120 dpf (Fig. 2B). In addition, *O. franksi* were not observed to spontaneously metamorphose even though they were maintained in the same culture conditions as *P. strigosa*. In order to emphasize the contrast in competence onset between *P. strigosa* and *O. franksi* we modeled two non-overlapping competence windows: 3 to 20 dpf for the short-PLD model (*P. strigosa* – like) and 20 to 120 dpf for the long-PLD model (*O. franksi* – like).

**Figure 2.**
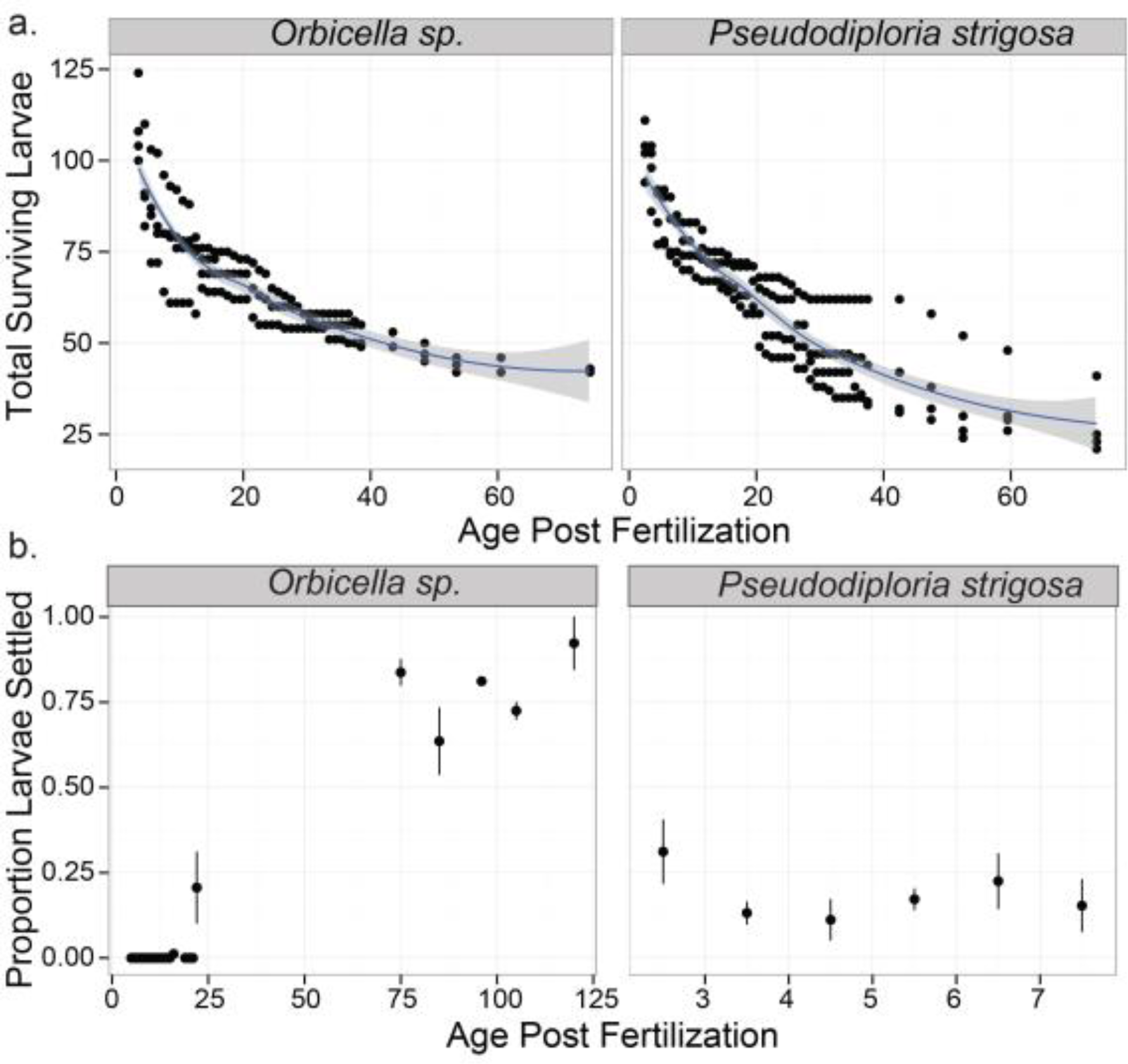
Coral larval physiological measurements: A. Mortality estimates for larvae from *Orbicella faveolata* and *P. strigosa* and across four culture replicates. Blue line is loess smoothing, grey shading indicates 95% confidence interval. B. Competency patterns observed for *Orbicella_franksi* and *P. strigosa* estimated by the mean proportion of larvae +/− SE settling in response to settlement cue over time. Note the differences in time scale for the two species.

### Short PLD results in occasionally abundant but highly variable reseeding

Short PLD occasionally lead to a very high probability of reseeding, but only on specific days in years when currents are favorable. In 2009, only larvae released from WFGB on August 10 and 11 could reseed either bank (probabilities of reseeding: 4.3E-03 and 1.3E-04, respectively), and the probability of reseeding on August 10 when released from WFGB was larger than any other release date in our analysis (Fig 3A,B, Table 1, Supplemental Fig S1). In 2010 and 2012 no short-PLD larvae were found within the boundaries of the FGB (Fig. 3D,H). In 2011, short-PLD larvae released from EFGB could reseed the FGB (simulation for WFGB for that year was not generated), but the probability was lower (5.4E-04) than on August 10 2009 (Fig. 3F, Table 1). The mean probability for short PLD larvae to reseed the FGB in 2009-2012 was 5.8E-04. The total reproductive output for *P. strigosa* at WFGB and EFGB was estimated at 4.7E+09 and 9.2E+09 eggs, respectively (Supplemental Table 1). Therefore, the mean potential reseeding events for short PLD larvae released in mass-spawn nights across all simulations run for 2009-2012 was 2.9E+06 (Fig. 4A). Notably, none of the simulated spawning events for the short PLD larvae resulted in any export to other reef systems besides the FGB.

**Figure 3.**
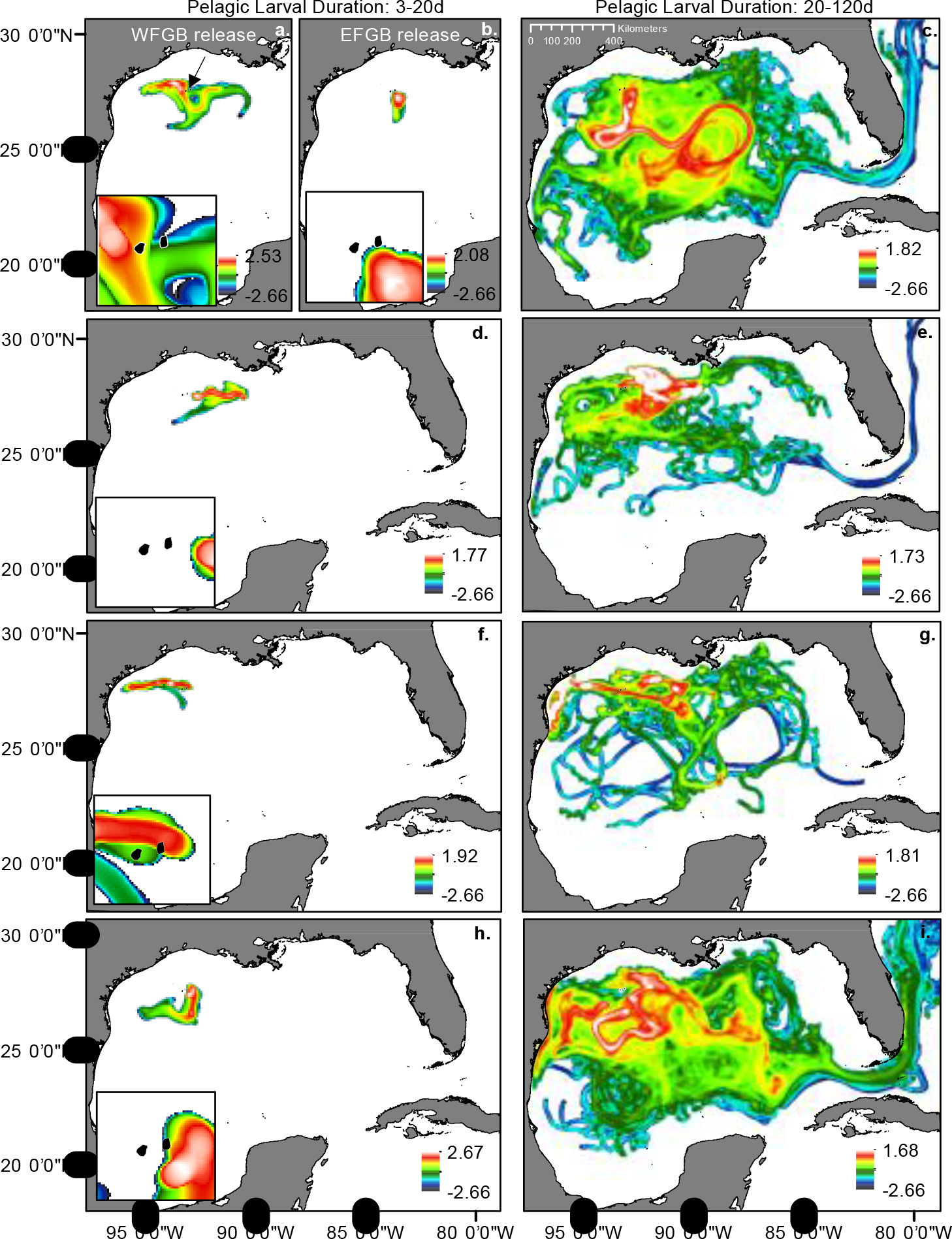
Surface heatmaps of particle dispersal from the Flower Garden Banks (FGB, arrow). Images show the density of particles for a 12-km radius around a 1-km cell integrated over pelagic larval durations (PLDs) of 3-20 days (A, B, D, F, H) and 20-120 days (C, E, G, I). Particle density is displayed on a log10 scale. Release dates are August 2009 (A, B, C), August 31, 2010 (D, E), August 19, 2011 (F, G) and August 8, 2012 (H, I). (A) Represents particles released from WFGB, (B-I) represent larvae released from EFGB. For short PLDs (3-20 days, A, B, D, F, H), insets show variation in particle’s reseeding area of FGB (black area) between years.

**Table 1.** Probability of larval reseeding to FGB and probability of larval export to other reefs. Proportion calculated as number of particles within area/total particles for set PLD (short: 3-20 days; long: 20-120 days).

| Date | Bank | PLD | Reproductive Output (# eggs) | P(reseeding) | P(export) | Total larvae reseeded | Total larvae exported |
| --- | --- | --- | --- | --- | --- | --- | --- |
| 10-Aug-09 | West | short | 4.73E+09 | 4.3 E-03 | 0 | 2.0E+07 | 0 |
|  |  | long | 1.02E+10 | 7.2E-05 | 4.1E-05 | 7.3E+5 | 4.2E+5 |
|  | East | short | 9.15E+09 | 0 | 0 | 0 | 0 |
|  |  | long | 2.36E+10 | 2.7E-04 | 4.4E-05 | 6.4E+6 | 1.0E+6 |
| 11-Aug-09 | West | short | 4.73E+09 | 1.3E-04 | 0 | 6.1E+5 | 0 |
|  |  | long | 1.02E+10 | 2.7E-04 | 2.0E-05 | 2.8E+6 | 2.0E+5 |
|  | East | short | 9.15E+09 | 0 | 0 | 0 | 0 |
|  |  | long | 2.36E+10 | 1.7E-04 | 5.7E-05 | 4.0E+6 | 1.3E+6 |
| 12-Aug-09 | West | short | 4.73E+09 | 0 | 0 | 0 | 0 |
|  |  | long | 1.02E+10 | 4.3E-05 | 1.1E-06 | 4.4E+5 | 1.1E+4 |
|  | East | short | 9.15E+09 | 0 | 0 | 0 | 0 |
|  |  | long | 2.36E+10 | 7.0E-05 | 1.3E-05 | 1.7E+6 | 3.1E+5 |
| 13-Aug-09 | West | short | 4.73E+09 | 0 | 0 | 0 | 0 |
|  |  | long | 1.02E+10 | 9.6E-05 | 0 | 9.8E+5 | 0 |
|  | East | short | 9.15E+09 | 0 | 0 | 0 | 0 |
|  |  | long | 2.36E+10 | 2.0E-05 | 5.6E-05 | 4.7E+5 | 1.3E+6 |
| 30-Aug-10 | East | short | 9.15E+09 | 0 | 0 | 0 | 0 |
|  |  | long | 2.36E+10 | 2.0E-03 | 3.1E-06 | 4.7E+7 | 7.3E+4 |
| 31-Aug-10 | East | short | 9.15E+09 | 0 | 0 | 0 | 0 |
|  |  | long | 2.36E+10 | 3.0E-05 | 0 | 7.1E+5 | 0 |
| 19-Aug-11 | East | short | 9.15E+09 | 5.4E-04 | 0 | 4.9E+6 | 0 |
|  |  | long | 2.36E+10 | 2.5E-04 | 0 | 5.9E+6 | 0 |
| 8-Aug-12 | West | short | 4.73E+09 | 0 | 0 | 0 | 0 |
|  |  | long | 1.02E+10 | 1.2E-04 | 1.2E-04 | 1.2E+6 | 1.2E+6 |
|  | East | short | 9.15E+09 | 0 | 0 | 0 | 0 |
|  |  | long | 2.36E+10 | 1.7E-06 | 4.0E-04 | 4.0E+4 | 9.4E+6 |
| 9-Aug-12 | West | short | 4.73E+09 | 0 | 0 | 0 | 0 |
|  |  | long | 1.02E+10 | 2.0E-04 | 5.5E-05 | 2.0E+6 | 5.6E+5 |
|  | East | short | 9.15E+09 | 0 | 0 | 0 | 0 |
|  |  | long | 2.36E+10 | 2.0E-05 | 2.4E-04 | 4.7E+5 | 5.7E+6 |

### Long PLD results in low but consistent reseeding

Simulations demonstrated that long-PLD larvae (competency from 20 to 120 dpf had similar average probabilities of reseeding (3.3E-04) as short-PLD larvae (5.8E-04), but the long-PLD larvae reseeded more consistently over the years (Table 1, Fig 3). On August 11 2009, the major spawning night for that year, the probabilities of long-PLD reseeding when released from WFGB or EFGB were 2.7E-04 and 1.7E-04, respectively (Fig. 3C, Table 1). On peripheral spawning nights in 2009 (August 10, 12 and 13), simulations showed similar probabilities of reseeding (Table 1). The highest probability of long PLD larval reseeding, 3eE-03, was observed for the August 30, 2010 simulation for the EFGB (Table 1, Supplemental Fig 3). Interestingly, larvae with longer PLDs had a dramatically higher probability of reseeding their home reef than larvae with short PLDs when released on August 30, 2010 (Table 1, Supplemental Fig 3). In 2011 and 2012, the probability of long PLD larvae reseeding the FGB ranged from 1.7E-06 to 1.2E-04 (Fig. 3G,I, Table 1). Total *Orbicella franksi* reproductive output was calculated as 1.02E+10 and 2.26E+10 eggs, for WFGB and EFGB respectively (Supplemental Table 1). At the FGB, *O. franksi* has much higher overall coral cover (26.90% at WFGB and 27.56% at EFGB) than *P. strigosa* (9.60% at WFGB and 8.20% at EFGB)^17^. These differences in percent cover account for the ~3-5x greater reproductive output calculated for *Orbicella sp*. when compared to *P. strigosa*. Therefore, for a mass spawn night, the average number of potential reseeding events of *Orbicella* larvae to the FGB between 2009 and 2012 was 8.3E+06 (Fig 4).

**Figure 4.**
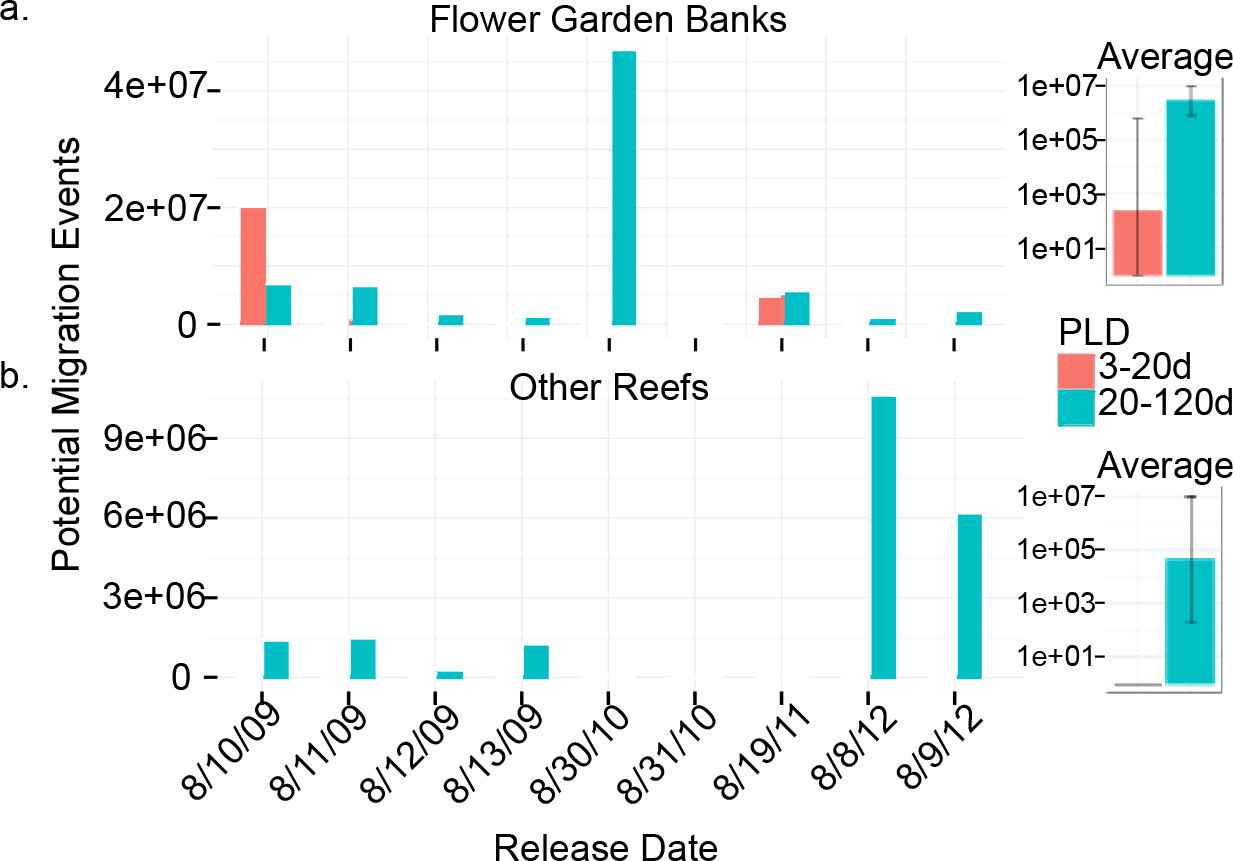
Larval particles with potential to colonize coral reef environments. Output from West and East FGB is summed. (A) Flower Garden Banks (reseeding) (B) Other reefs as defined in [UNEP-WCMC, WorldFish Centre, WRI, TNC]. PLD = pelagic larval duration. Insets show harmonic mean number of particles for each PLD +/− SD.

### Long PLD enables larval export beyond the FGB

No export events were observed for short PLD larvae, but larvae with PLDs of 20-120 days remained in the water column long enough to be exported to reefs outside of the FGB. For 2009 simulations, the probability of larval export to other reefs ranged from
1. 1E-06 to 5.7E-05, similar to the probability of reseeding back to the FGB. After accounting for total *O. franksi* reproductive output, this resulted in 1.1E+04-1.3E+06 potential larval export events in 2009, depending on the day and bank of release (Table 1, Fig. 4B). In 2009, long PLD larvae were observed in Broward and Palm Beach, the Florida Keys (Dry Tortugas up to Elliott Key), Bay of Campeche reefs (Bajos del Norte, Bajo Madagascar, Alacranes, Arenas, Triangulos, Lobos, Bajo Madaga) (reefs defined in Sanvicente-Añorve et al 2014 and Millennium Coral Reef Mapping Project 2005) and Northern Cuba (west of Cayo Coco). This larval export varied greatly from year to year: in 2010 only a single larva in the simulation (released from WFGB on August 30) dispersed to a reef other than FGB (Little Bahama Banks), resulting in an export probability of 3.1E-06, and no larvae were exported to other reefs in 2011. In 2012 simulations suggested export to other reefs at even higher rates than those observed in the 2009 simulation: probability of larval export ranged from 5.5E-05 to 1.20E-04. After considering reproductive output, between 5.6E+05 and 9.4E+06 possible larvae were exported in 2012 (Table 1, Fig 4B). The destinations of exported larvae in 2012 were Broward and Palm Beach, the middle to upper Florida Keys (Islamorada up to Elliott Key), Bay of Campeche reefs (Bajos del Norte, Alacranes, Arenas) and the Bahamas (Cay Sal Banks and Little Bahama Banks).

## Discussion

### Corals from the FGB exhibit unusual competency dynamics

Larvae of most broadcast-spawning scleractinian corals become competent upon completion of larval development, typically 4-6 dpf, after which competence declines^18– 20^. Both FGB coral species studied here, *Pseudodiploria strigosa* and *Orbicella franksi*, deviate from this pattern, but in opposing directions. *P. strigosa* becomes competent by 2.5 dpf, while *O. franksi* cultured in identical conditions, did not exhibit competence until 20 dpf. In addition, once competence was acquired in *O. franksi*, no decline in competence was detected, even up to 120 dpf. Davies et al (2014) observed similar competency onset ranges: 3-4 dpf for *Pseudodiploria* (formerly *Diploria) strigosa* and 14-20 dpf for *O. franksi*, indicating that competency patterns reported here are consistent across years for populations of these species at the FGB. While two-day pre-competency periods have been observed in *Pectinia lactuca* and *Platygyra sinensis* from Singapore^21^ and *Platygyra daedlia* on the Great Barrier Reef^18^ no other study has ever reported delay of competency by as much as 20 dpf, as was observed here in FGB *O. franksi*. Although there is no published data on competence in *O. franksi* from other regions, larvae of a congener, *Orbicella faveolata*, from Belize became competent at a typical 6 dpf^22^. Similarly, *O. faveolata* larvae from the Florida Keys began gaining competence at 6 dpf and exhibited 100% competence by 24 dpf^23^. It is therefore possible that unusual competence dynamics of FGB corals may be specific to the FGB populations-the possibility of which requires clarifications in future studies.

### PLD and reseeding potential

Along the lines of Darwin’s hypothesis about the loss of flight ability in insects and birds following colonization of remote islands^24^, one might predict that corals at an isolated reef such as the FGB would be selected for short PLDs to facilitate reseeding. However, we find that both short-PLD and long-PLD models result in similar average FGB reseeding probabilities (Fig. 4), suggesting that FGB reseeding might not necessarily require the loss of long-distance dispersal potential. The possibility that the optimal PLD for the FGB reseeding could be bimodal has been suggested by an earlier study tracking drifters released from the FGB during two annual coral spawning events in 1997 & 1998^25^. There, it was demonstrated that while there was a high likelihood of drifters’ returning to the FGB within the first 30 days, they were also entrained in Loop Current eddies and could recirculate back to the FGB after several months. The two coral species modeled here appear to take advantage of these two alternative reseeding strategies, but whether their unusual PLDs are a result of local selection at the FGB remains to be determined, especially considering that these species are rarely observed to recruit^26^.

A large body of research links short PLD to high reseeding and low connectivity, while long PLD is linked to lower reseeding values and higher connectivity^7,27,28^. Given that our long-PLD species had a late onset of competency and was able to both reseed and disperse, our results support the alternative view that this paradigm may be too simplistic within the real seascape^29^. Still, our study agrees with previous genetic^30^ and oceanographic modeling work performed in the region^25,30^ in that we find that species with early onset of competency and short PLDs are capable of reseeding, but are indeed highly isolated from neighboring reefs.

### Dispersal is highly variable among larval cohorts

Larval release timing significantly affected dispersal probabilities (Fig. 3). In 2010 the probability of FGB reseeding was far greater for long PLD larvae than for short PLD larvae and only one long PLD larva was exported to another reef. In contrast, in 2011 simulations, similar probabilities of reseeding were observed for both short and long PLD larvae, with no larval export to other reefs regardless of PLD. Finally, in 2009 and 2012 long PLD larvae had high probabilities of export while short PLD larvae exhibited the highest probability of reseeding in 2009 and no chance of reseeding in 2012.

Spatial variation in the pattern of larval release also interacts with temporal variation, resulting in dramatically different dispersal patterns, even over relatively small geographic distances^31^. The east and west FGB are only 18 kilometers apart, but in 2009, EFGB short PLD larvae dispersed further east and had no chance of reseeding across all four days while WFGB short PLD larvae drifted west but were more likely to be maintained in the vicinity of FGB or circulate back (Fig. 3AB, Supplemental Fig S1). The opposite pattern was observed in 2012 where only EFGB larvae reseeded while WFGB larvae drifted west and were never able to return to FGB. However, in spite of this variance, recurrence of high-probability reseeding events indicates that FGB coral populations are likely to be demographically self-sustaining.

### Long PLD is essential for larval export from the FGB to other reefs

Surface currents in the GOM tend to be dominated by the Loop Current (LC), which is a continuation of the Caribbean current that intrudes the GOM through the Yucatan Channel ^32^. The position and degree of LC intrusion into the GOM is variable in season and across years ^33^ and can vary from flowing directly into the Florida current to intruding the GOM as far as 29.1°N (http://oceancurrents.rsmas.miami.edu/caribbean/loop-current.html). The degree of LC intrusion into the GOM influences the likelihood of large warm water eddies being cast off and flowing westward into the GOM. These eddies can be large enough that a full rotation can be up to 30 days^34^. Large scale eddy formation from the LC is irregular^33^, but is more prevalent in the summer months^35^ when broadcast-spawning corals release their gametes. Lugo-Fernández *et al*. (2001) suggest that as much as 43% of larvae released from FGB are likely to get caught up in these offshore eddies. The results of our simulations support this assertion: we find that larval transport in the GOM is highly affected by LC eddy circulation. For example, in 2009, high numbers of larvae became entrained in a large eddy that was detached from the LC on Sept 2^36^. The density of larvae released from the FGB in other years appeared to be less affected by these large-scale eddies. However, in nearly all of our simulations with long PLD larvae (20-120 days), surviving particles were eventually able to enter the LC, likely through eddy entrainment, and be dispersed by either the Florida Current (moving towards the Florida Keys and Miami or the Bahamas) or westward through the Yucatan Current and potentially into the Campeche shelf reefs.

Interestingly, larvae from the FGB were never observed to enter the Western Caribbean directly, presumably due to the LC acting as a dispersal barrier (Fig 3). Connectivity between the FGB and the Western Caribbean must therefore be facilitated by stepping-stones, which likely include Florida, Bahamas and potentially Cuba. Although our modeling data suggest that some FGB larvae can disperse to reefs in the southern GOM, other research has shown that these reefs on the Campeche banks are likely sink populations due to the constraint of the LC on dispersal into the Western Caribbean^37^. Johnson *et al*. (2013) modeled red snapper larval dispersal from the southern GOM and reported very high probabilities of reseeding (67-73%) with 0.33% of larvae arriving at other reefs, including the FGB, but no larvae dispersed outside the GOM. Thus, the FGB might be an important stepping-stone between the southern GOM and other reefs in the Northern Caribbean.

### FGB as a refugium

The ability of FGB to act as a refugium is contingent on the population’s ability to withstand stress and the frequency of disturbances. In other potential coral refugia sites in the southern hemisphere, bleaching events have been reported^39,40^. However, despite the occurrence of these events at other high latitude sites, the FGB continues to be resilient to these disturbances and remains one of the healthiest reefs in the Caribbean^3^. Our data show that for species with long PLDs, the FGB can act as a source of larvae for distant reefs in the southern GOM, Florida, the northern and western Bahamas and Northern Cuba, highlighting the potential of the comparatively remote and pristine FGB to act as a refugium. Our simulations predict the possibility for large export events across the Florida reef tract, which are orders of magnitude higher than previously predicted^25^. This result demonstrates that highly detailed models including specific times and locations of larval release as well as important life history traits, such as larval competency parameters, can drastically change predictions of larval transport between sites and overall source/sink dynamics.

Characterizing potential refugia populations is critical for reef management and the design of reserve networks, as individuals migrating from these sites could modulate characteristics of future populations^29,41^. Both the FGB and the Florida Keys are maintained as United States national marine sanctuaries, however, anthropogenic influences and coral cover of endangered groups, including *Orbicella*, is dramatically different between these ecosystems (Galindo *et al*. 2006; Palandro *et al*. 2008, Emma Hickerson, personal communication). Fishing and tourism strongly affect Florida reef ecosystems and hard coral cover has significantly declined over the last 30 years^43^. Our data demonstrate the potential for larvae released from the FGB to contribute to *Orbicella franksi* populations in the Florida Keys. Continued protection of highly fecund colonies from the pristine FGB reefs may be important for maintaining larval supply and genetic diversity along degraded Florida reefs.

### Outlook for future research

The largest remaining knowledge gap in the modeling of coral larval dispersal is how to translate the probability of larvae arriving to a certain location (such as the results of our modeling) into the actual recruitment rate. Throughout this paper we followed other authors in implicitly assuming (i) that recruitment probability is directly proportional to the larval arrival probability, (ii) that this proportion is the same for different coral species, and (iii) that this proportion is independent of the environmental conditions at the target location. One indication that these assumptions might be unrealistic is the size-frequency distribution of adult corals observed at the FGB (Fig. 1B): *P. strigosa* shows a significantly higher proportion of smaller colonies than *O. franksi*, which, since the growth rates of the two species are similar^44^, suggests higher recent recruitment of *P. strigosa*. However, our model predicts that short-PLD larvae of *P. strigosa* should be on average less likely to arrive to FGB than the long-PLD larvae of *O. franksi* (Fig. 4A). One explanation for this apparent discrepancy is that *P. strigosa* larvae might be more efficient at recruiting, or suffer less post-settlement mortality than *O. franksi*. It is also conceivable that the recruitment probability might scale non-linearly with numbers of arriving larvae such that recruitment effectively occurs only when very high numbers of larvae arrive, which would be more likely for the short-PLD *P. strigosa* (Fig. 4A). Finally, the strength of ecological barriers to larval dispersal (i.e., due to environmental differences rather than physical separation of the habitats) remains entirely unknown. It is possible that larvae produced in one type of habitat would not be physiologically and/or genetically predisposed to survive in a different type of habitat, the situation termed “phenotype-environment mismatch”^45^. This mismatch could be particularly relevant for FGB-originating larvae, since the FGB is a very unusual reef compared to the rest of the Caribbean. More research is needed to investigate these possibilities and develop more realistic models.

## Methods

### Study species

*Pseudodiploria strigosa* was used both for larval mortality and competency trials, but for logistical reasons two different species of the *Orbicella* species complex were used to measure larval traits: *Orbicella faveolata* for mortality and *O. franksi* for competency. Due to negligible differences in mortality rates across genera (*P. strigosa* versus *O. faveolata)* we assumed that their average larval mortality rates would be a good approximation for both *Pseudodiploria* and the *Orbicella* species complex.

### Size-frequency transects

On August 8 and 9, 2012 divers completed a total of eight size frequency transects targeting *Orbicella franksi* and *Pseudodiploria strigosa* following protocols established by the Florida Reef Resiliency Program^46^ (http://frrp.org). Four transects were completed at both the WFGB (27° 52.526 N,-93° 48.836 W) and EFGB (27° 54.516 N,-93° 35.831 W) (Fig 1). In brief, communities were surveyed using 10-m^2^ belt transects, randomly placed at each site. The diameter, height and percent live tissue of all corals ≥4 cm in diameter were recorded. Colony sizes were log-transformed to visualize size-frequency differences between the two corals and these size-frequency distributions were compared using a Wilcoxon rank sum test in R.

### Larval rearing

Samples were collected under the FGBNMS permit # FGBNMS-2009-005-A3. On the evening of August 18, 2011 (eight days after the full moon, 2115CDT), gamete bundles from ≥3 individuals of each broadcast-spawning Caribbean coral species (*P. strigosa* and *O. franksi)* were collected from the east FGB. Gamete bundles were brought to the surface and allowed to cross-fertilize in 3L of 1 μm filtered seawater (FSW) for one hour in sterile 6L plastic containers. Excess sperm were removed by rinsing through 150μm nylon mesh. Larvae were reared in 1 μm FSW in three replicate plastic culture vessels stocked at a density of 2 larvae per ml in a temperature controlled room (28°C). Larvae were transferred to the laboratory at the University of Texas at Austin on August 21, 2011 and used in all competency trials.

On the evening of August 8, 2012 (2330CDT) divers collected gamete bundles from four spawning *O. faveaolata* colonies and the next evening (August 9, 2012 at 2115CDT) divers collected from eight spawning *P. strigosa*. Cultures for both species were fertilized and maintained as described in 2011. Larvae were transferred to the University of Texas at Austin on August 10, 2012 and these larvae were used in mortality trials. All research in 2012 was completed under the Flower Garden Banks National Marine Sanctuary permit #FGBNMS-2012-002.

### Mortality trials

Mortality trials began on August 11, 2012 for *O. faveolata* and *P. strigosa*. Four mortality trials per species were conducted. Each trial started with 100 larvae housed in a bowl with 0.5 L of FSW. For the first 38 days the surviving swimming larvae were counted daily and transferred into new bowls. After that the larvae were counted every five days until 74 days post fertilization (dpf).

### Competency trials

For competency trials each well in a six-well plate received 10 ml of FSW and a drop of a uniform slurry of finely ground crustose coralline algae collected at the F GB, which has previously been shown to elicit settlement in both species^47^. Twenty larvae of *O. franksi* or *P. strigosa* were then added to each well (n = 3 replicates per species) and the proportions of metamorphosed larvae (visual presence of septa) were quantified after 24 h using a stereomicroscope MZ-FL-III (Leica, Bannockburn, IL, USA). Trials began 2.5 dpf and were repeated until all larvae that remained in vessels had spontaneously metamorphosed.

### Biophysical model

We used the Conn4D biophysical dispersal model ^48^ to estimate dispersal patterns for each species independently. Conn4D uses oceanographic current information in conjunction with an advection diffusion scheme and individual-based behaviour to simulate larval trajectories. Oceanographic data was derived from the HYbrid isopycnal Coordinate Model (HYCOM)^49^. We parameterized the models to reflect the exponential
decrease in larval mortality measured for both genera. Simulations did not encompass a full 4D model since coral larvae were assumed to stay in surface waters as passive drifters. Therefore, the depth of particles was kept at a constant 5 meters. Dates of particle release were as follows: August 10^th^, 11^th^, 12^th^ and 13^th^ 2009, August 30^th^ and 31^st^ 2010, August 11^th^ 2011 and August 8^th^ and 9^th^ 2012. These dates were the dates of observed spawning at FGB, eight days after the full moon in late July/August (Emma Hickerson, personal communication). Coral spawning at the FGB is highly predictable^50^, however the night of the mass spawn can be flanked by nights that have smaller outputs of gametes. Therefore for some years we ran simulations on these flanking days. Simulated particles were released from either West (27.83° N 93.83° W) or East (28.00° N 93.58° W) FGB. For each simulation, 1000 particles were released and allowed to drift for 120 days. Output from each simulation was recorded as a text file with latitude, longitude, depth, time, distance and duration traveled for each particle, as in^48^. Output files were split into two non-overlapping time windows of 3-20 days for the “short PLD” and 20-120 days for the “long PLD” datasets.

### Analysis

Simulated particle dispersal was visualized in ArcGIS 10.2. Input data was transformed using the ‘Project (Data Management)’ tool from Decimal Degrees to a Projected Coordinate system NAD 1983 UTM Zone 16N and visualized within the Gulf of Mexico. The location of the particles in the Gulf of Mexico was summarized as a density surface by calculating the number of particles on or around a 12-km radius and raster output was set to a 1-km cell size. Density surface rasters were log10 transformed using the ‘raster calculator’. The number of total particles within the boundary of the FGBNMS and within the boundary of other reefs (as defined by UNEP-WCMC, WorldFish Centre, WRI, TNC) for each PLD was calculated using the ‘select by location’ tool. This gives the total number of times any particle intersects the specified boundary, not the total number of particles with a final destination within the boundary. In some cases, a particle stays within the specified boundary for several days, which influences its probability of being a migrant within the specified reef boundary. For each simulation, the number of particles within each boundary during a specified PLD was divided by the total number of particles for that simulation, giving a probability of particle existence within each specified boundary. This probability of particle presence within the FGB and in other reefs was multiplied by the total reproductive output of each species from each bank to give a value representing the potential migration events. Total reproductive output for East and West FGB was estimated by multiplying fecundity data (number of eggs/m^2^) from parameters measured in Szmant (1986) for *Pseudodiploria strigosa* (35,200 eggs/m^2^) and in Szmant *et al*. (1997) for *O. franksi* (27,000 eggs/m^2^) by area of coral cover of each species at each bank^2,17^ (Supplemental Table 1).

## Author Contributions

SWD and JTK conceived the study. Data was collected by SWD and CDK. Data was analyzed by SWD and MES. All authors wrote and reviewed the manuscript.

## Acknowledgements

We are grateful to the staff and volunteers at the Flower Garden Banks National Marine Sanctuary (FGBNMS) for providing boat time, field assistance and permits across several coral spawning field seasons. Thanks to Eli Meyer for assistance in the field. We acknowledge Thanapat Pongwarin and Sarah Guermond for their assistance in the laboratory measuring life history traits. In addition, we acknowledge ARCCoE for the opportunity to initiate this collaborative study. This research was funded in part by the PADI Foundation Grant to S.W.D. Computing resources were provided by Australia’s National Computational Infrastructure (NCI). JK publishes with the permission of the Chief Executive Officer of Geoscience Australia.

---

**Supplementary Figure 1:**
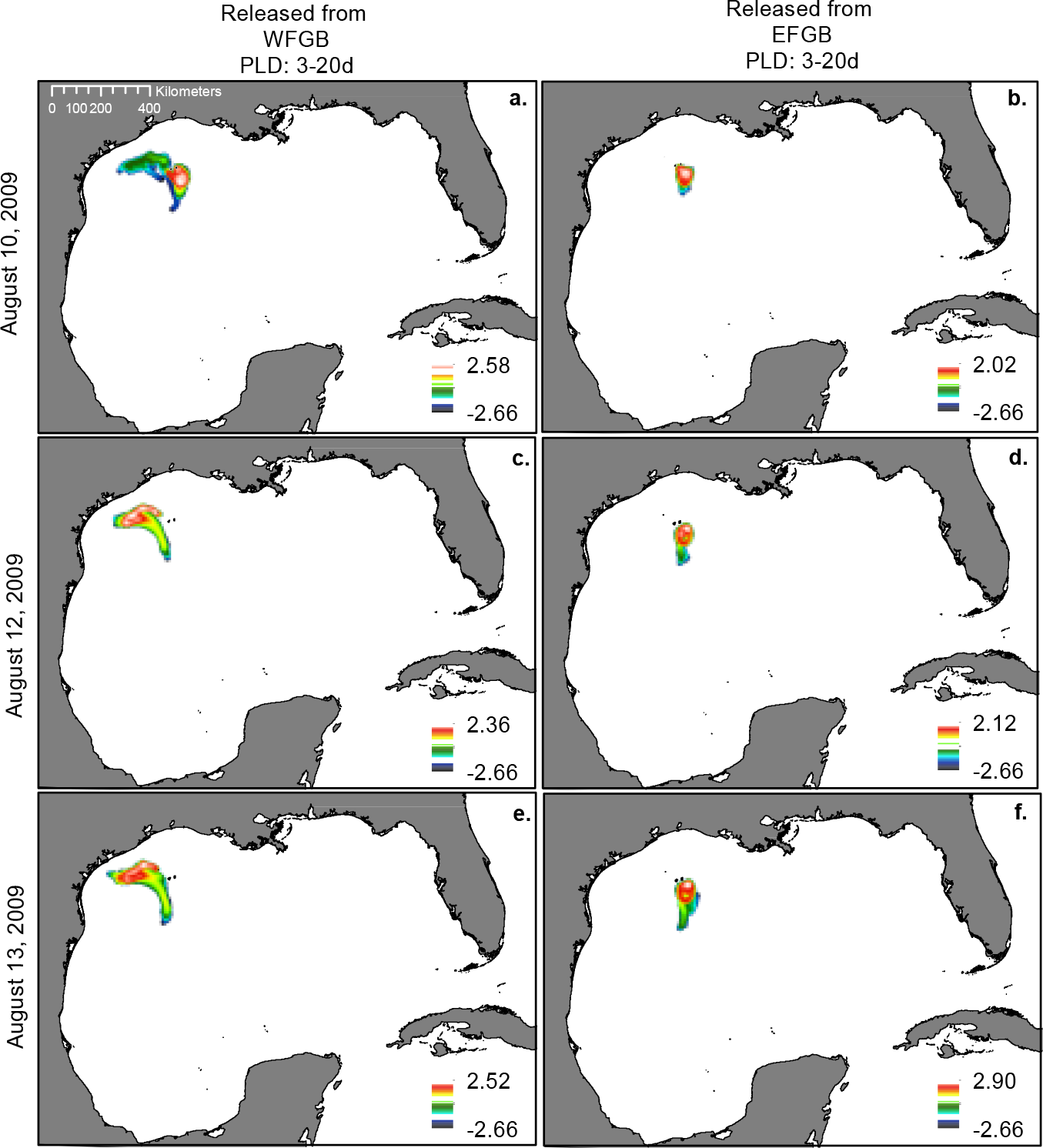
Surface heatmaps of larval particle dispersal from West (A, C, E) and East (B, D, F) Flower Garden Banks, TX in 2009. Images show the density of larval particles for a 12-km radius around a 1-km cell, integrated over 3-20 days after release. Particle density displayed on a log10 scale. (A-B) particles were released on August 10, 2009. (C-D) particles were released on August 12, 2009. (E-F) particles were released on August 13, 2009.

**Supplementary Figure 2:**
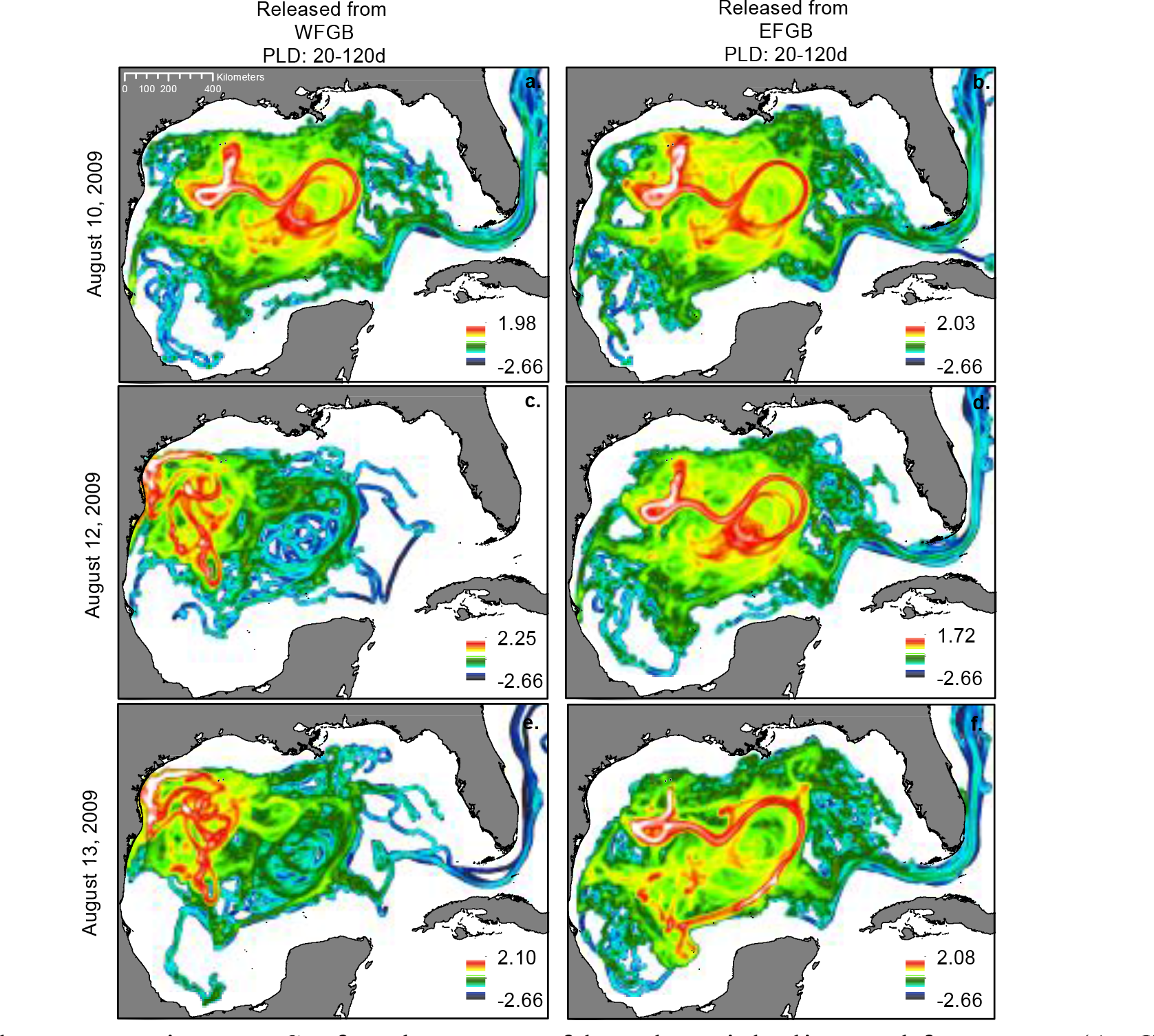
Surface heatmaps of larval particle dispersal from West (A, C, E) and East (B, D, F) Flower Garden Banks, TX in 2009. Images show the density of larval particles for a 12-km radius around a 1-km cell, integrated over 20-120 days after release. Particle density displayed on a log10 scale. (A-B) particles were released on August 10, 2009. (C-D) particles were released on August 12, 2009. (E-F) particles were released on August 13, 2009.

**Supplementary Figure 3:**
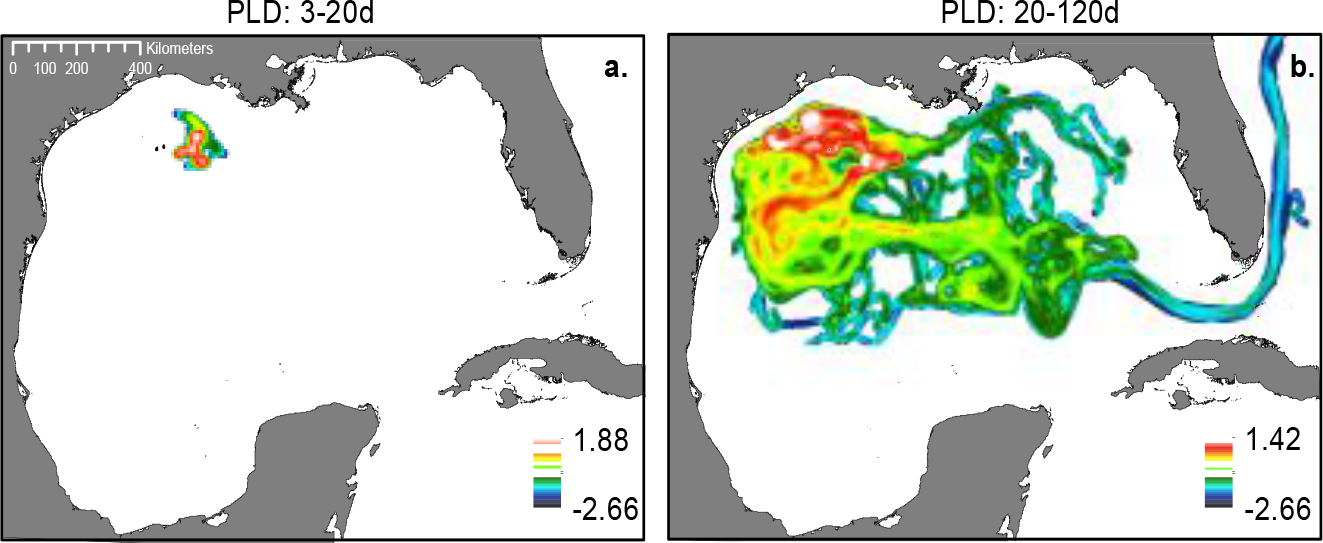
Surface heatmaps of larvel particle dispersal from East Flower Garden Banks on August 30, 2010. Images show the density of larval particles for a 12km radius around a 1-km cell, integrated over 3-20 days (A) and 20-120 days (B) after release. Particle density displayed on a log10 scale.

**Supplementary Figure 4:**
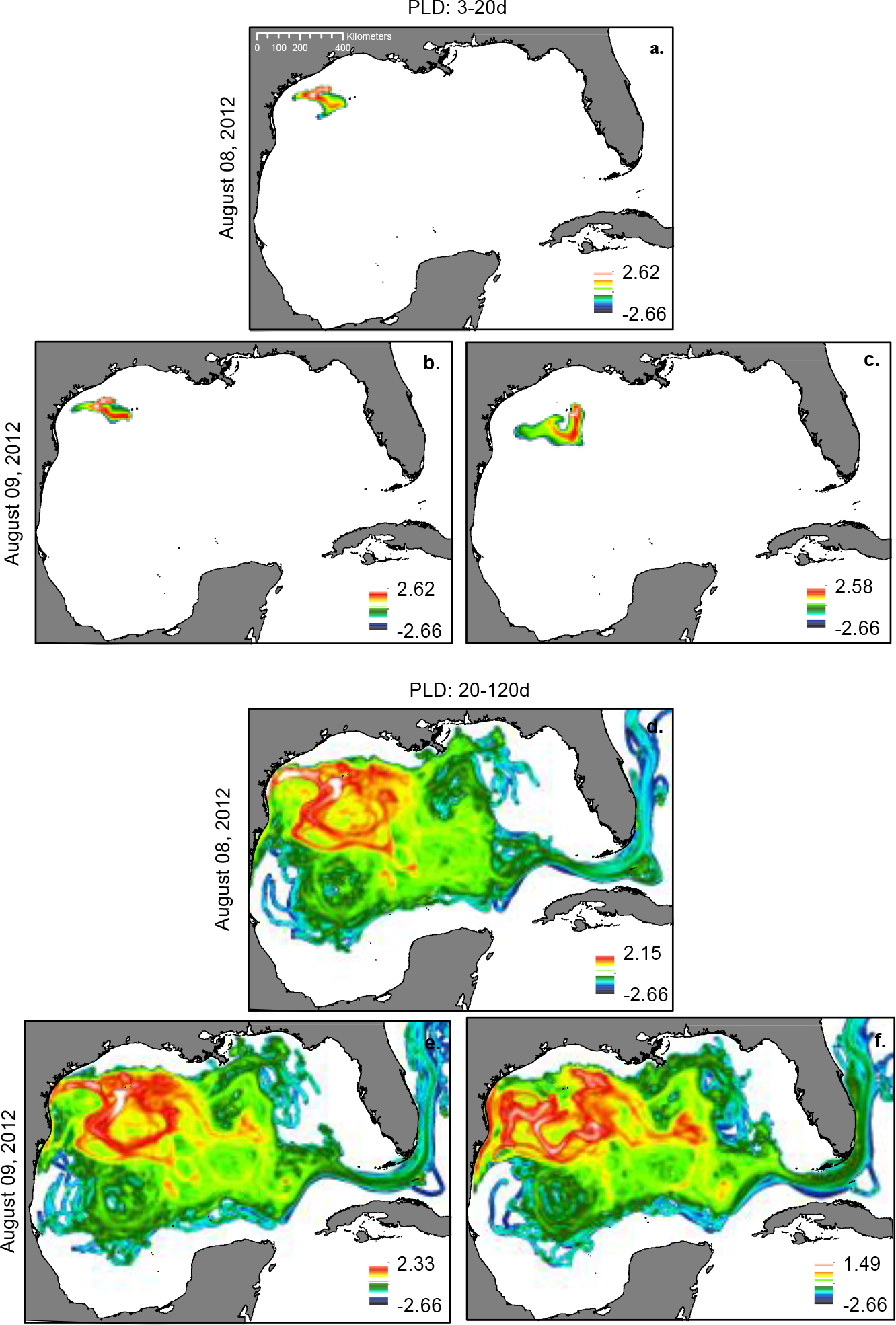
Surface heatmap of larval particle dispersal from West (A, C, D, E) and East (C, E) Flower Garden Banks, TX in 2012. Images show the density of larval particles for a 12-km radius around a 1-km cell, integrated over 3-20 days (A, B, C) and 20-120 days (D, E, F) on August 8 (A-C) and August 9 (D-E) 2012.

**Supplemental Table 1:** Reproductive output estimated for both *Pseudodiploria strigosa* and *Orbicella franksi* for both the East and West Flower Garden Banks. Total area of WFGB reef cap <150ft deep is 1.4E+6 m^2^ and total area of EFGB reef cap <150ft deep is 3.17E+6 m^2 17^. Eggs per polyp were estimated at 270/cm^2^ (90 per polyp × 3 polyps/cm^2^) for *Orbicella franksi* and 352/cm^2^ (8 eggs/gonad × 22 gonads/polyp × 2 polyps/cm^2^) for *Pseudodiploria strigosa*.

|  | % Cover | Area (m<sup>2</sup>) | Eggs/m<sup>2</sup> | Total eggs |
| --- | --- | --- | --- | --- |
| WFGB |  |  |  |  |
| *P. strigosa* | 9.60 | 1.34E+05 | 3.52E+04 | 4.73E+09 |
| *O. franksi* | 26.90 | 3.77E+05 | 2.70E+04 | 1.02E+10 |
| EFGB |  |  |  |  |
| *P. strigosa* | 8.2 | 2.60E+05 | 3.52E+04 | 9.15E+09 |
| *O. franksi* | 27.56 | 8.74E+05 | 2.70E+04 | 2.36E+10 |

## References

1. Gardner, T. a, Côté, I.M., Gill, J. a, Grant, A. & Watkinson, A.R. Long-term region-wide declines in Caribbean corals. Science 301, 958–60 (2003).

2. Schmahl, G.P., Hickerson, E.L. & Precht, W.F. in Coral Reefs of the USA (eds. Riegl, B.M. & Dodge, R.E.) 221–261 (Springer Netherlands, 2008). doi:10.1007/978-1-4020-6847-8_6

3. Zimmer, B., Duncan, L., Aronson, R.B., Deslarzes, K. J. P. & Dies, D.R. Longterm monitoring at the East and West Flower Garden Banks, 2004-2008. Volume I: Technical Report. (US Dept of the Interior, Bureau of Ocean Energy Management, Regulation, and Enforcement, Gulf of Mexico OCS Region, 2010).

4. Rezak, R., Gittings, S.R. & Bright, T.J. Biotic assemblages and ecological controls on reefs and banks of the Northwest Gulf of Mexico. Am. Zool. 30, 23–35 (1990).

5. Sale, P.F. in Population and Community Biology 197–210 (1991).

6. Kinlan, B.P. & Gaines, S.D. Propagule Dispersal in Marine and Terrestrial Environments: a Community Perspective. Ecology 84, 2007–2020 (2003).

7. Shanks, A.L., Grantham, B.a. & Carr, M.H. Propagule Dispersal Distance and the Size and Spacing of Marine Reserves. Ecol. Appl. 13, 159–169 (2003).

8. Davies, S., Treml, E., Kenkel, C. & Matz, M. Exploring the role of Micronesian islands in the maintenance of coral genetic diversity in the Pacific Ocean. Mol. Ecol. 24, 70–82 (2015).

9. Jones, G., Srinivasan, M. & Almany, G. Population Connectivity and Conservation of Marine Biodiversity. Oceanography 20, 100–111 (2007).

10. Gaines, S.D., Gaylord, B., Gerber, L.R., Hastings, A. & Kinlan, B.P. The Ecological Consequences of Dispersal in the Sea. Mar. Popul. Connect. 20, 90–99 (2007).

11. Van Oppen, M. J. H., Peplow, L.M., Kininmonth, S. & Berkelmans, R. Historical and contemporary factors shape the population genetic structure of the broadcast spawning coral, Acropora millepora, on the Great Barrier Reef. Mol. Ecol. 20, 4899–4914 (2011).

12. Cowen, R.K. in Ecology of Coral Reef Fishes: Recent Advances 149–170 (Academic press, 2002).

13. Cowen, R.K. Population Connectivity in Marine Systems. Oceanography 20, 14–21 (2007).

14. Munday, P.L. et al. Climate change and coral reef connectivity. Coral Reefs 28, 379–395 (2009).

15. Kendall, M.S., Poti, M. & Karnauskas, K.B. Climate change and larval-transport in the ocean: Fractional effects from physical and physiological factors. Glob. Chang. Biol. 1532–1547 (2015). doi:10.1111/gcb.13159

16. Wilson, L.J. et al. Climate-driven changes to ocean circulation and their inferred impacts on marine dispersal patterns. Glob. Ecol. Biogeogr. 1–17 (2016). doi:10.1111/geb.12456

17. Johnston, M.A., Nuttall, M.F., Eckert, R.J. & Embesi, J.A. Long-Term Monitoring at the East and West Flower Garden Banks, 2013 Annual Report. (2014).

18. Miller, K. & Mundy, C. Rapid settlement in broadcast spawning corals: implications for larval dispersal. Coral Reefs 22, 99–106 (2003).

19. Harrison, P. & Wallace, C. in Coral reef ecosystems 133–207 (Elsevier, 1990).

20. Baird, A.H. The ecology of coral larvae: settlement patterns, habitat selection and the length of the larval phase. (James Cook University, 2001).

21. Tay, Y.C., Guest, J.R., Chou, L.M. & Todd, P.A. Vertical distribution and settlement competencies in broadcast spawning coral larvae: Implications for dispersal models. J. Exp. Mar. Bio. Ecol. 409, 324–330 (2011).

22. Ritson-Williams, R., Arnold, S.N., Paul, V. J. & Steneck, R.S. Larval settlement preferences of Acropora palmata and Montastraea faveolata in response to diverse red algae. Coral Reefs 33, 59–66 (2014).

23. Vermeij, M., Fogarty, N. & Miller, M. Pelagic conditions affect larval behavior, survival, and settlement patterns in the Caribbean coral Montastraea faveolata. Mar. Ecol. Prog. Ser. 310, 119–128 (2006).

24. Darwin, C. On the origin of species by means of natural selection, or the preservation of favoured races in the struggle for life. A facsimile of the first edition with an introduction by Ernst Mayr. (Harvard University Press., 1859).

25. Lugo-Fernández, A., Deslarzes, K. J. P., Price, J.M., Boland, G.S. & Morin, M. V. Inferring probable dispersal of Flower Garden Banks coral larvae (Gulf of Mexico) using observed and simulated drifter trajectories. Cont. Shelf Res. 21, 47–67 (2001).

26. Davies, S.W., Matz, M.V. & Vize, P.D. Ecological complexity of coral recruitment processes: effects of invertebrate herbivores on coral recruitment and growth depends upon substratum properties and coral species. PLoS One 8, e72830 (2013).

27. Sponaugle, S. et al. Predicting Self-Recruitment in Marine Populations: Biophysical Correlates and Mechanisms. Bull. Mar. Sci. 70, 341–375 (2002).

28. Foster, N.L. et al. Connectivity of Caribbean coral populations: complementary insights from empirical and modelled gene flow. Mol. Ecol. 21, 1143–57 (2012).

29. Cowen, R.K. & Sponaugle, S. Larval dispersal and marine population connectivity. Ann. Rev. Mar. Sci. 1, 443–466 (2009).

30. Galindo, H.M., Olson, D.B. & Palumbi, S.R. Seascape Genetics: A Coupled Oceanographic-Genetic Model Predicts Population Structure of Caribbean Corals. Curr. Biol. 16, 1622–1626 (2006).

31. Kough, A.S. & Paris, C.B. The influence of spawning periodicity on population connectivity. Coral Reefs 34, 753–757 (2015).

32. Oey, L., Ezer, T. & Lee, H. Loop Current, rings and related circulation in the Gulf of Mexico: A review of numerical models and future challenges. Circ. Gulf Mex. Obs. Model. 161, 31–56 (2005).

33. Alvera-Azcárate, A., Barth, A. & Weisberg, R.H. The Surface Circulation of the Caribbean Sea and the Gulf of Mexico as Inferred from Satellite Altimetry. J. Phys. Oceanogr. 39, 640–657 (2009).

34. Berger, T.J. Louisiana-Texas Shelf Physical oceanography Program Eddy Circulation Study. I, 1992–1995 (1997).

35. Chang, Y.L. & Oey, L.Y. Why does the Loop Current tend to shed more eddies in summer and winter? Geophys. Res. Lett. 39, 1–7 (2012).

36. Taylor, P., Lindo-atichati, D., Bringas, F. & Goni, G. International Journal of Remote Loop Current excursions and ring detachments during 1993-2009. 37–41 (2013).

37. Sanvicente-Añorve, L., Zavala-Hidalgo, J., Allende-Arandía, M. & Hermoso-Salazar, M. Connectivity patterns among coral reef systems in the southern Gulf of Mexico. Mar. Ecol. Prog. Ser. 498, 27–41 (2014).

38. Johnson, D.R., Perry, H.M. & Lyczkowski-Shultz, J. Connections between Campeche Bank and Red Snapper Populations in the Gulf of Mexico via Modeled Larval Transport. Trans. Am. Fish. Soc. 142, 50–58 (2013).

39. Harrison, P.L., Dalton, S.J. & Carroll, A.G. Extensive coral bleaching on the world’s southernmost coral reef at Lord Howe Island, Australia. Coral Reefs 30, 775 (2011).

40. Thomson, D.P., Bearham, D. & Graham, F. High latitude, deeper water coral bleaching at Rottnest Island, Western Australia. Coral Reefs 30, 1107 (2011).

41. Palumbi, S.R. Population genetics, demographic connectivity, and the design of marine reserves. Ecol. Appl. 13, 146–158 (2003).

42. Palandro, D.A. et al. Quantification of two decades of shallow-water coral reef habitat decline in the Florida Keys National Marine Sanctuary using Landsat data (1984-2002). Remote Sens. Environ. 112, 3388–3399 (2008).

43. Donahue, S. et al. The State of Coral Reef Ecosystems of the Florida Keys. The state of coral reef ecosystems of the United States and Pacific Freely Associated States: 2008. (eds Waddell JE, Clarke AM) (2005). at http://ccma.nos.noaa.gov/ecosystems/coralreef/coral2008/pdf/FloridaKeys.pdf

44. Muslic, A. et al. Linear Extension Rates of Massive Corals from the Dry Tortugas National Park (DRTO), Florida. (2013).

45. Marshall, D.J., Monro, K., Bode, M., Keough, M.J. & Swearer, S. Phenotype-environment mismatches reduce connectivity in the sea. Ecol. Lett. 13, 128–40 (2010).

46. Wagner, D.E., Kramer, P. & Woesik, R. Van. Species composition, habitat, and water quality influence coral bleaching in southern Florida. Mar. Ecol. Prog. Ser. 408, 65–78 (2010).

47. Davies, S.W., Meyer, E., Guermond, S.M. & Matz, M.V. A cross-ocean comparison of responses to settlement cues in reef-building corals. PeerJ 1–20 (2014). doi:10.7717/peerj.333

48. Kool, J.T. & Nichol, S.L. Four-dimensional connectivity modelling with application to Australia’s north and northwest marine environments. Environ. Model. Softw. 65, 67–78 (2015).

49. Chassignet, E. P. et al. The HYCOM (HYbrid Coordinate Ocean Model) data assimilative system. J. Mar. Syst. 65, 60–83 (2007).

50. Vize, P.D., Embesi, J.a., Nickell, M., Brown, D.P. & Hagman, D.K. Tight temporal consistency of coral mass spawning at the Flower Garden Banks, Gulf of Mexico, from 1997-2003. Gulf Mex. Sci. 23, 107–114 (2005).

51. Szmant, A.M. Reproductive ecology of Caribbean reef corals. Coral Reefs 5, 43–53 (1986).

52. Szmant, A. M., Weil, E., Miller, M.W. & Colon, D.E. Hybridization within the species complex of the scleractinan coral Montastraea annularis. Mar. Biol. 129, 561–572 (1997).

